# Identification of a Third Period-tuning Site in Cyanobacterial Clock Protein KaiC

**DOI:** 10.64898/2026.05.11.724173

**Authors:** Kota Horiuchi, Yoshihiko Furuike, Kumiko Ito-Miwa, Yasuhiro Onoue, Shuji Akiyama

## Abstract

KaiC, a clock protein in cyanobacteria, cycles between dephosphorylated and phosphorylated states in a 24-hour period in the presence of KaiA and KaiB. We identified the 322nd residue of KaiC as a third example of period-tuning sites. 322nd–site-directed saturation mutagenesis resulted in a variety of KaiC mutants exhibiting either shortened or lengthened cycles. The tunable range of the periods was from approximately 11 to 78 h without significantly compromising temperature compensation. We conducted biochemical analyses of the 322nd variants and examined their predicted structural models. In contrast to another known period-tuning site, where the period decreases sharply as the side-chain volume increases due to mutations, the cycle lengths correlate only modestly with bulkiness at the 322nd residues. The 322nd residue is located in a C-terminal domain of KaiC and influences ATPase cycles in both the C-terminal domain and an N-terminal domain through its interaction with a flexible loop connecting the two domains. The structural models predict that placing less bulky but polar side chains, such as serine and threonine, at the 322nd position leads to the formation of a hydrogen-bonding network between that site and the loop. This reduces the mobility of the loop, resulting in the longer cycles due to decreases in the ATPase activity of the N-terminal domain. Conversely, placing bulky residues such as phenylalanine at the 322nd position appears to alter the loop structure, shortening the periods by enhancing the ATP activities of both the domains. The third period-tuning mechanism is distinct from other known mechanisms.

**Significance Statement:** A Kai-protein clock system serves as a model for studying how long circadian rhythms are achieved. We identified the 322nd residue of KaiC as a third example of period-tuning sites that allow tuning of the period in either long- and short-period directions. The third period-tuning mechanism differs from the two previously known types in several respects. Previous studies have suggested that the ATPase activity in an N-terminal domain of KaiC is the primary regulator of the period. On the other hand, the 322nd residues of KaiC can affect the period by activating the ATPase cycle in its C-terminal domain. Our findings will stimulate future studies on the period-tuning mechanism mediated by the ATPase activity in the C-terminal domain of KaiC.

## Introduction

Circadian clocks are endogenous systems that rhythmically control a variety of biological processes of organisms with a period of approximately 24 h (1). The circadian clocks share three common characteristics. First, circadian rhythms are autonomous and persist even under constant conditions. Second, the period length is temperature compensated. Third, the phase of the circadian clock shifts forward or backward in response to external stimuli, synchronizing with that of environmental cycles.

A question, regarding the first property, of how such the long cycle of 24 hours is achieved and regulated remains unresolved in the field of chronobiology. Since the first isolation of single-gene mutations that alter the period length (2), a variety of clock-related proteins and their mutants have been identified (3). According to accumulated results to date, multiple amino acids affecting period length are widely distributed in clock proteins, and the structural motifs and domains contributed by these amino acids are also diverse.

One such example is a clock protein PERIOD (PER) (2, 4). PER contains multiple scattered amino acids that receive post-translational phosphorylation modification (phosphorylation sites), and even substituting just one of these can alter the period length (5, 6). At the same time, point mutations in the interaction interfaces where PER binds to other clock proteins also results in the period changes (7, 8). A tau mutation in casein kinase 1 (CK1) disrupts it anion binding, shortening the period by shifting the substrate specificity of CK1 toward phosphodegron site (6). A similar trend has been reported for a clock protein CRYPTCHROME-1 (CRY1), in which the period-altering sites are identified widely around its co-factor binding pocket comprised of a phosphate-binding loop (P-loop) and a C-terminal lid domain (9). In this way, the amino acids affecting the period length are diverse both in position and structure, spanning phosphorylation sites (10-13), ligand-binding sites (9, 14), inter-domain interfaces within the same clock protein (15, 16), and inter-molecular interaction interfaces (6-8, 17). This diversity is one of the factors making it difficult to establish a unified picture regarding period length determination.

“Period-tuning (PT) site” is a key concept for addressing such the issue. The PT site refers to a specific amino acid that allows tuning of the period length in either long- and short-period directions depending on the type of post-substituted residues. The bidirectionality suggests that the PT sites possess a more direct causal relationship with the period-determining mechanism in the clock proteins. However, most of the above examples exhibit unidirectional changes in the cycle length or lack validation of the bidirectionality. There are only a limited number of the known PT sites: the 662nd position in PER2 (18), the 243rd position in CRY1 (9), and two PT sites in KaiC, a core clock protein from *Synechococcus elongatus* PCC 7942 (19).

KaiC consists of 519 amino acids and possesses an N-terminal (CI) domain and a C-terminal (CII) domain. Six subunits of KaiC are self-assembled into a double-ring hexamer (Fig. 1A), in which every CI–CI and CII–CII interface binds one molecule of adenosine triphosphate (ATP) or adenosine diphosphate (ADP) (20, 21). In the presence of KaiA, KaiB, and ATP, S431 and T432 in the CII domain are rhythmically phosphorylated and then dephosphorylated (P-cycle) with the circadian period (22-25).

**Figure 1.**
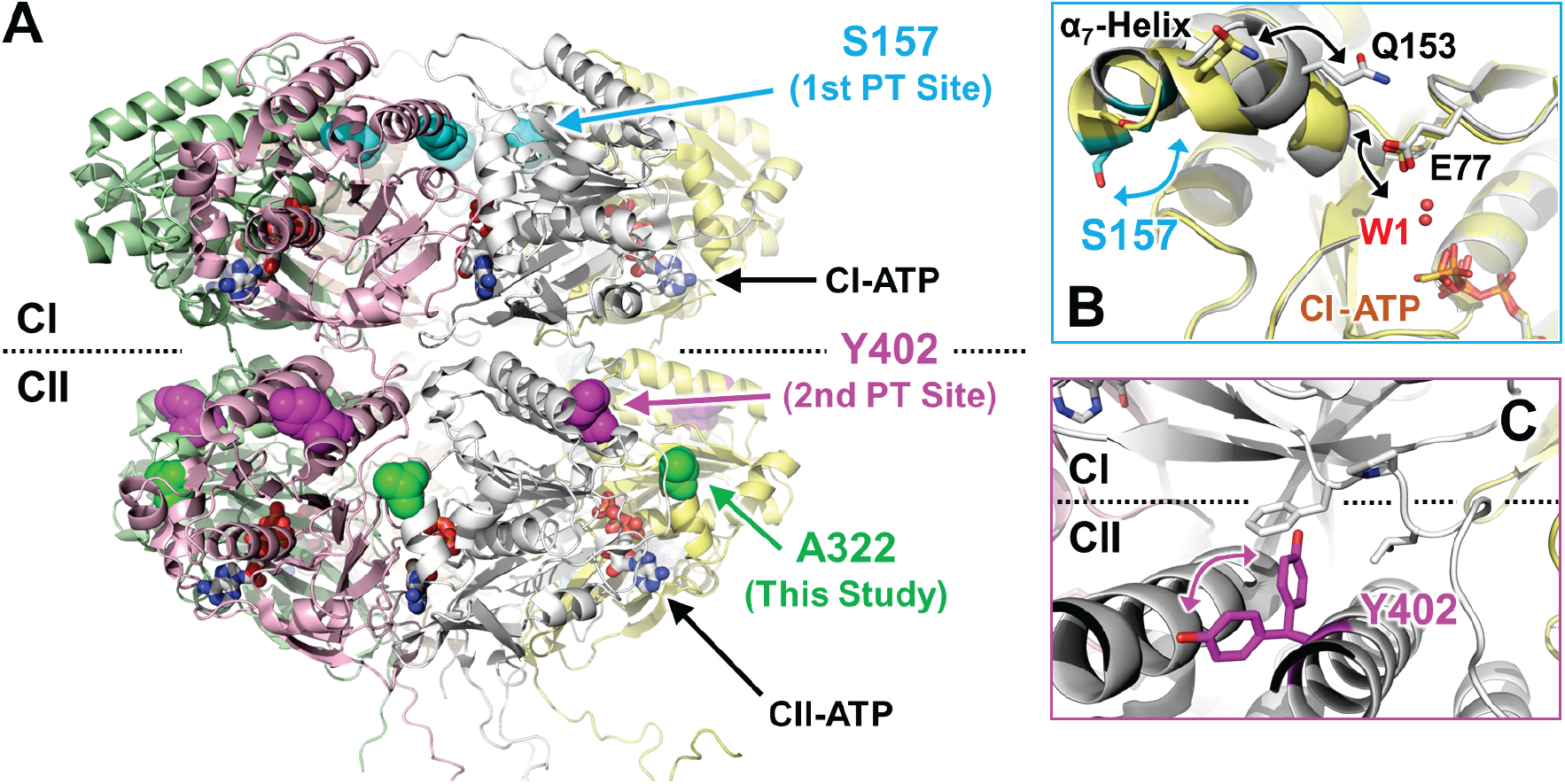
Period-tuning (PT) sites in KaiC. (**A**) Two known and one newly identified PT sites mapped on the crystal structure of KaiC (PDB ID: 2GBL). (**B**) The 157th position as the first PT site (19, 26). A positional shift of a lytic water molecule (W1) is coupled to a transition of the α_7_-helix between a frayed (gray) and a capped (yellow) structure via Q153 and E77 (PDB ID: 4TL7 and 4TLC). The S157C and S157P replacements result in destabilization and stabilization, respectively, of the capping structure of the α_7_-helix (26). (**C**) The 402nd position as the second PT site (28) located near a contact region between the CI and CII domains (PDB ID: 7S65) (29).

The first identified PT site is the 157th residue in the CI domain (Fig. 1A), which corresponds to a Ser residue in wild-type KaiC (KaiC^WT^). While a KaiC mutant with the S157P substitution (KaiC^S157P^) exhibits a shorter P-cycle (21 h) than KaiC^WT^ (24 h), KaiC^S157C^ results in a longer P-cycle (25 h) (19, 26). The P-cycle frequency correlates positively with the ATPase activity of the CI domain (27), and that the PT mechanism at the 157th position is based on this correlation (26). As shown in Fig. 1B, the 157th residue plays a role in capping the N-terminus of the α_7_-helix located nearby CI-ATP. The S157C substitution biases the α_7_-helix into a frayed conformation, in which a lytic water molecule (W1) is away from CI-ATP and sequestered at a position unfavorable for the ATP hydrolysis (Fig. 1B). On the other hand, the S157P substitution stabilizes a capped conformation and achieves the higher ATPase activity by positioning W1 closer to CI-ATP in that conformation (Fig. 1B).

The 402nd residue is identified as the second PT site harboring a conserved Tyr residue (28). Although the 402nd residue is classified within the CII domain according to the primary sequence, it is actually located near a contact region between the CI and CII domains (Fig. 1A). The smaller the side-chain volume at the 402nd residue, the more the period is lengthened: KaiC^Y402W^ and KaiC^Y402A^ show the P-cycle lengths of 15 and 158 h, respectively (28). A later cryoelectron microscopy study (29) suggests that the 402nd residue can affect the communication between the CI and CII domains (Fig. 1C).

Here we report the 322nd residue as the third PT site. This PT site possesses a conserved Ala residue and is classified within the CII domain in terms of both the primary sequence and three-dimensional structure (Fig. 1A). A series of 322nd mutants exhibited dramatical changes of the cycle length both *in vivo* and *in vitro* in both the long- and short-period directions. Despite the wide-ranging changes, the cycle lengths and ATP consumption rates are temperature compensated in most of the 322nd mutants. Although the 322nd residue is located near the CII–CII interface (Fig. 1A), its side-chain volume showed only a modest correlation with the cycle length, in sharp contrast to the second PT site. The third PT mechanism, by which the 322nd residue modulates the cycle length by affecting the dual ATPase sites in the CI and CII domains, appears to be distinct from the first and second PT mechanisms.

## Results

### 322nd residue serving as the third PT site

*In vitro* P-cycles of KaiC^WT^ and 19 types of A322 variants were measured in the presence of KaiA and KaiB at 30°C. The cycle length of KaiC^WT^ was 23.4 ± 0.9 h at 30°C (A322A in Fig. 2), which was consistent with the results of previous studies (22-24). Among a series of the variants examined (Fig. 2, *SI Appendiex*, Fig. S1), a mutant with the A322T substitution (KaiC^A322T^) exhibited the longest period (78.3 ± 1.1 h), while KaiC^A322F^ had the shortest cycle length (10.9 ± 0.1 h). Longer periods were confirmed for a total of seven substitutions: A322T (78.3 ± 1.1 h), A322P (41.2 ± 2.2 h), A322R (27.5 ± 1.0 h), A322S (27.1 ± 0.2 h), A322N (25.7 ± 1.2 h), A322C (25.2 ± 0.8 h), and A322K (24.8 ± 0.5 h). A total of nine substitutions resulted in shorter periods in descending order: A322D (23.0 ± 0.3 h), A322Q (22.1 ± 0.7 h), A322E (21.2 ± 0.2 h), A322M (20.8 ± 0.3 h), A322H (18.0 ± 0.4 h), A322W (15.9 ± 0.4 h), A322L (13.8 ± 0.1 h), A322Y (12.9 ± 0.1 h), and A322F (10.9 ± 0.1 h). Although the remining three variants, KaiC^A322V^, KaiC^A322I^, and KaiC^A322G^, were arrhythmic under the *in vitro* condition at 30°C, the bidirectional changes in the cycle period were observed in more than 80% of the 322nd variants. These results clearly suggest that the 322nd residue potentially functions *in vitro* as a PT site.

**Figure 2.**
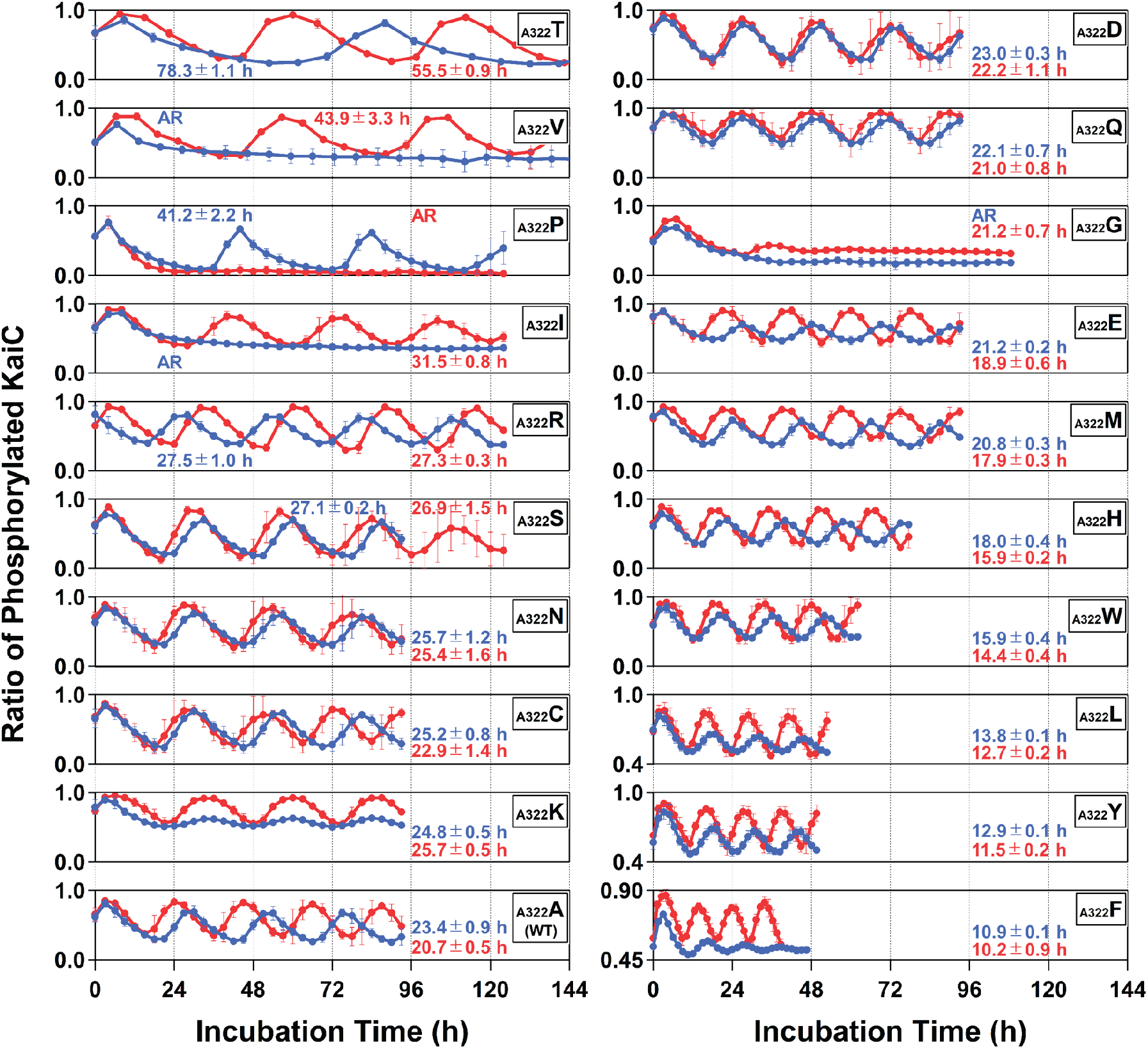
*In vitro* phosphorylation cycles of KaiC^WT^ and a series of its mutants with substitutions at the 322nd position. Blue and red circles correspond to the means ± s.d. from independent experiments conducted at 30 and 40°C, respectively, on different days (see *SI Appendix*, Fig. S1 and S3 for individual data). Note that mutants with lower amplitudes (KaiC^A322L^, KaiC^A322Y^, and KaiC^A322F^) are plotted using adjusted vertical scale ranges for clarity of presentation. To achieve the reproducibility of rhythms, KaiC^A322S^ and KaiC^A322V^ at 40°C required 1.5-fold (0.06 mg/ml) higher KaiA concentrations.

To test whether this also applies under *in vivo* conditions, we observed bioluminescence rhythms of cyanobacterial reporter strains carrying a series of the A322 substitutions in KaiC. At 30°C, a wild-type strain (A322A in Fig. 3) showed a robust bioluminescent rhythm with a period of 24.9 h. The A322T substitution exhibited the longest period also *in vivo* (59.1 h) as well as *in vitro*. Unfortunately, we were unable to quantify the period of the A322F strain possibly due to its extremely low amplitude. However, the A322Y substitution, which resulted in the second shortest period *in vitro* (Fig. 2), showed the shortest cycle length *in vivo* (22.0 h) (Fig. 3). The *in vivo* periods of other mutants varied bidirectionally (Fig. 3, *SI Appendeix*, Fig. S2) mostly in accordance with the relationship confirmed *in vitro* (Fig. 2). Consequently, the cycle lengths under *in vitro* and *in vivo* conditions showed a good correlation at 30°C (Fig. 4A), with a few exceptions of the arrhythmic cases mentioned above. The 322nd residue of KaiC is the PT site that functions both *in vitro* and *in vivo*.

**Figure 3.**
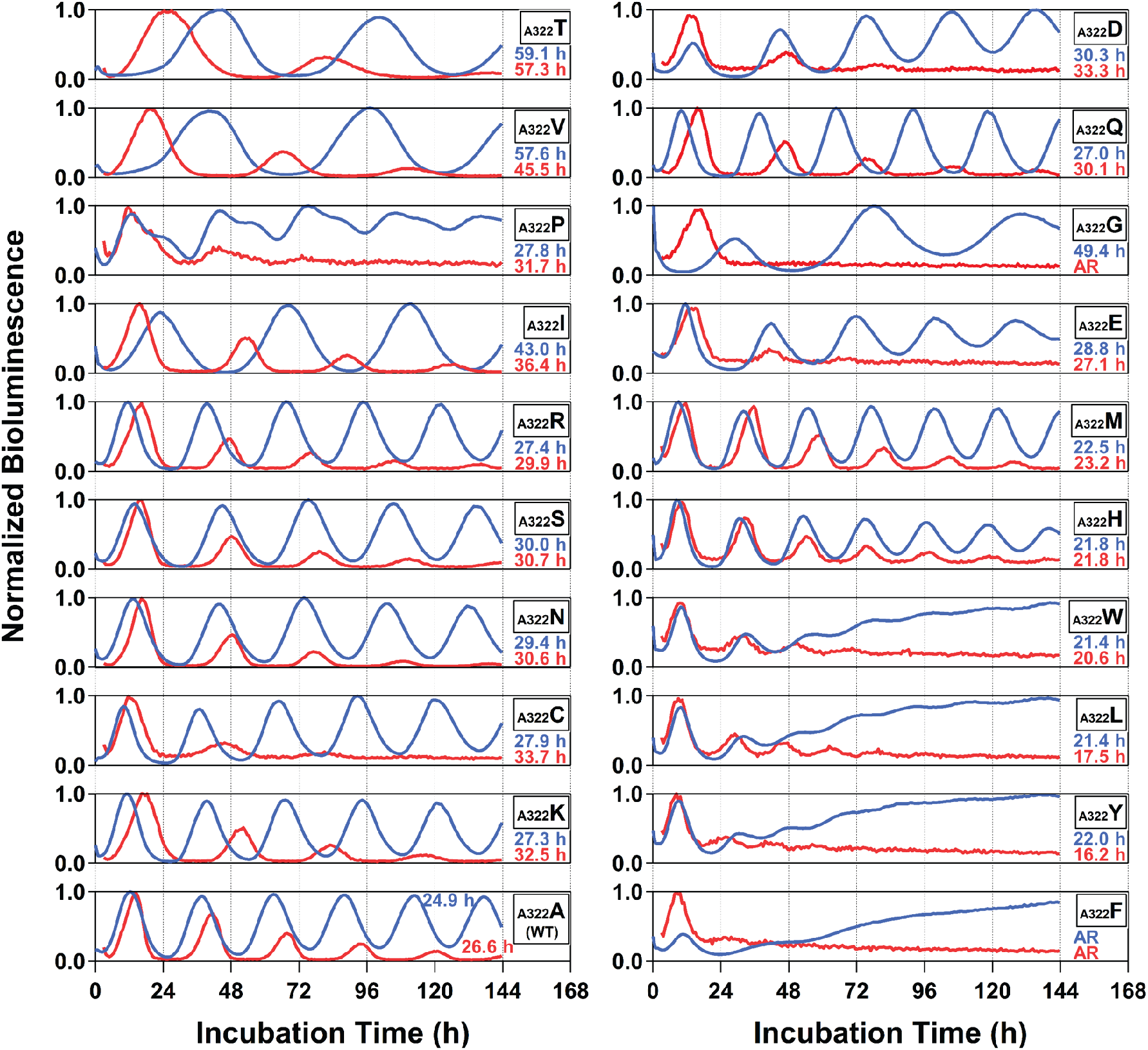
*In vivo* bioluminescence rhythms of *Synechococcus elongatus* PCC 7942 reporter strains carrying *kaiC*^*WT*^ or a series of its mutants with substitutions at the 322nd position under conditions of constant light. Blue and red lines correspond to the experiments conducted at 30 and 40°C, respectively (see *SI Appendix*, Fig. S2 and S4 for individual data).

**Figure 4.**
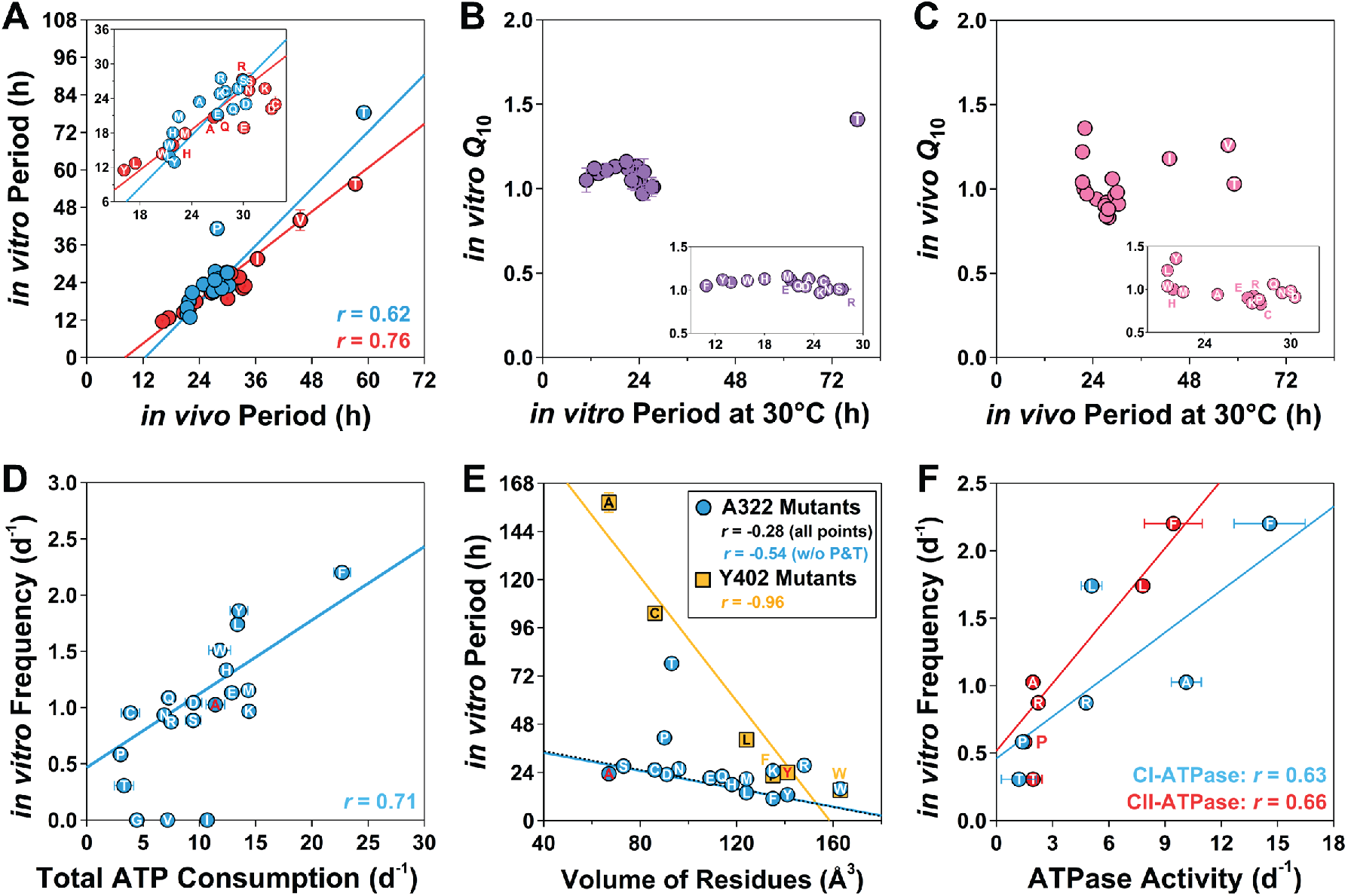
Periods, temperature coefficients (*Q*_10_), and ATP consumption rates of 322nd variants. (**A**) Correlation between *in vitro* and *in vivo* periods at 30°C (blue) and 40°C (red). The single letter indicated at each data point represents the amino acid after substitution. For data points with small errors, error bars are hidden behind the markers. (**B**) *Q*_10_ of the *in vitro* periods. (**C**) *Q*_10_ of the *in vivo* periods. (**D**) *In vitro* P-cycle frequencies plotted against total ATP consumption rates at 30°C. Arrhythmic mutants such as KaiC^A322G^, KaiC^A322V^, and KaiC^A322I^ are plotted as having P-cycle frequencies of zero. Data points corresponding to KaiC^WT^ are shown in red single letters. (**E**) Dependences of the *in vitro* periods on volume of the 322nd (blue) and 402nd residues (orange). A blue line: correlation excluding the data points for KaiC^A322P^ and KaiC^A322T^. A black dotted line: correlation over all the data points. (**F**) *In vitro* P-cycle frequencies plotted against CI-ATPase (blue) and CII-ATPase activities (red) at 30°C.

To examine the effect of the 322nd substitutions on the temperature compensation, we measured *in vitro* and *in vivo* rhythms at 40°C (Fig. 2 and 3, *SI Appendix*, Fig. S3 and S4). Interestingly, the two mutants KaiC^A322V^ and KaiC^A322I^ were arrhythmic *in vitro* at 30°C, but exhibited robust P-cycles at 40°C (Fig. 2). In contrast, the phosphorylated fraction of KaiC^A322P^ transitioned from being rhythmic to arrhythmic as the temperature increased by 10°C. Except for such cases, the correlation between the *in vitro* and *in vivo* periods remained high even at 40°C (Fig. 4A). We calculated a temperature coefficient (*Q*_10_) using the cycle lengths at 30°C and 40°C. *Q*_10_ is a factor by which the reaction rate is accelerated by rising the temperature by 10°C, ranging from approximately 2 to 3 in ordinary enzymes (30). *In vitro Q*_10_ ranged from 0.97 ± 0.03 (KaiC^A322K^) to 1.41 ± 0.03 (KaiC^A322T^) with an average of 1.1 ± 0.1 (Fig. 4B), while *in vivo Q*_10_ ranged from 0.83 (KaiC^A322C^) to 1.36 (KaiC^A322Y^) with an average of 1.0 ± 0.1 (Fig. 4C). Moderate effects on *Q*_10_ were seen both *in vitro* and *in vivo*, especially in the long-period mutants. However, *in vitro Q*_10_ of KaiC^A322T^ and *in vivo Q*_10_ of KaiC^A322Y^ were well below 2–3, indicating that they still remain within the range of mild temperature compensation. The 322nd residue is the third PT site, which allows the bidirectional tuning of the cycle length without significantly compromising the temperature compensation (Fig. 4B and 4C) and the coupling between *in vitro* and *in vivo* rhythms (Fig. 4A).

### ATPase consumption rates of the third PT mutants

The majority of the total ATP consumption by KaiC^WT^ is attributable to its CI-ATPase cycle (26, 27, 31), whereas only a small fraction is consumed by its kinase/phosphatase (27) and ATPase activities (31) in the CII domain. A notable correlation is observed between the total ATP consumption rate of KaiC and the frequency of the P-cycle (26, 27), and this correlation persists even in the first (26) and second PT mutants (28). We investigated the total ATP consumption of the 322nd variants at 30°C and found that the correlation holds true also in the third PT mutants. As shown in Fig. 4D, the total ATP consumption rates of the low-frequency mutants KaiC^A322P^, KaiC^A322T^, and KaiC^A322C^ were much lower than that of KaiC^WT^, whereas the high-frequency variant KaiC^A322F^ exhibited the highest ATP consumption. With the exception of the three arrhythmic mutants (KaiC^A322G^, KaiC^A322V^, and KaiC^A322I^) plotted in Fig. 4D as having P-cycle frequencies of zero, the correlation coefficients at 30°C was estimated to be 0.71 for the remaining 322nd mutants. These results indicate that the total ATP consumption rate is one of the key factors determining the P-cycle frequency of the third PT variants.

We previously reported that the 402nd residue is the 2nd PT site that influences the CI-ATPase activity through the CI–CII contact region (20, 29), because the volume of the 402nd residue correlates with the ATP consumption rate and the P-cycle period (28). To examine whether the 322nd residue shares a similar PT mechanism with the 402nd residue, the P-cycle periods were plotted against the side-chain volume of the 322nd residues (Fig. 4E). The slope of a linear regression line for the 322nd residues was approximately five times less significant than that for the 402nd residues. This marked difference suggests that the PT mechanism at the 322nd residue is not the same as that of the 402nd residue. Consistent with this, the introduction of the Y402W substitution known to have the shortest-period among the 402nd substitutions further shortened the period of KaiC^A322L^ (*SI Appendix*, Fig. S5A) and KaiC^A322Y^ (*SI Appendix* Fig. S5B).

We found that the 322nd residue affects the ATP consumption rate in the CII active site (CII-ATP consumption). To estimate the CII-ATP consumption, we introduced the E77Q/E78Q substitutions to KaiC^WT^ and five types of the 322nd variants (KaiC^A322F^, KaiC^A322L^, KaiC^A322T^, KaiC^A322P^, and KaiC^A322R^), and inactivated their dual-catalytic glutamates in the CI domain (31, 32). The short-period 322nd mutants of KaiC^A322F^ and KaiC^A322L^ exhibited higher CII-ATP consumption rates than KaiC^WT^ and the long-period 322nd mutants of KaiC^A322T^, KaiC^A322P^, and KaiC^A322R^ (red bars in *SI Appendix*, Fig. S6). The CI-ATPase activities of these mutants were estimated by calculating the difference in the total ATP consumption rates between the samples with and without the E77Q/E78Q substitutions (orange bars in *SI Appendix*, Fig. S6).

The amount of ATP consumed as the kinase and phosphatase activities can be calculated using turnover rates of unsynchronized P-cycles of KaiC in the absence of KaiA and KaiB (*SI Appendix*, Fig. S7) (33). The ATP consumption as the kinase/phosphatase activities at 30°C were relatively low, and were estimated as 0.14, 0.08, 0.46, 0.25, 0.49, and 0.46 d^-1^ for KaiC^A322T^, KaiC^A322P^, KaiC^A322R^, KaiC^WT^, KaiC^A322L^, and KaiC^A322F^, respectively (green bars in *SI Appendix*, Fig. S6). Then, we estimated the CII-ATPase activities by subtracting the kinase/phosphatase activities from the CII-ATP consumption rates (blue bars in *SI Appendix*, Fig. S6). Interestingly, the P-cycle frequencies of the 322nd mutants correlated slightly better with the CII-ATPase activity (*r* = 0.66) than with the CI-ATPase activity (*r* = 0.63) (Fig. 4F). These results suggest that the 322nd residue tunes the P-cycle frequency by influencing both the CI-ATPase and CII-ATPase cycles.

## Discussion

Although KaiC^A322F/Y402W^ could not be expressed and purified due its instability, the 322nd mutants with the second- and third-shortest periods exhibited the even shorter cycle lengths after the introduction of the Y402W replacement (KaiC^A322L/Y402W^ and KaiC^A322Y/Y402W^ in *SI Appendix*, Fig. S5). The 322nd position affects the CI and CII active sites through a mechanism different from that of the second PT site.

In contrast to the 402nd position (28), the cycle length was not highly dependent on the volume at the 322nd positions (Fig. 4E). Even treating KaiC^A322T^ and KaiC^A322P^ as outliers, the spread of the data points in Fig. 4E was pronounced particularly for the mutants with large side-chain volumes at the 322nd position (*r* = −0.54); e.g., in KaiC^A322K^, KaiC^A322R^, and KaiC^A322W^. Although the side chains of Leu and Met are nearly the same in volume (Fig. 4E), they differ in terms of bulkiness due to their distinct branching patterns. In fact, the spread was reduced by replotting the periods of the 322nd variants with respect to bulkiness proposed by Zimmerman et al (34) (*r* = −0.62 in Fig. 5A). Although not as pronounced as in the 402nd variants (Fig. 4E), a clear trend emerged in which the cycle lengths tend to decrease moderately as bulkiness at the 322nd position increases.

**Figure 5.**
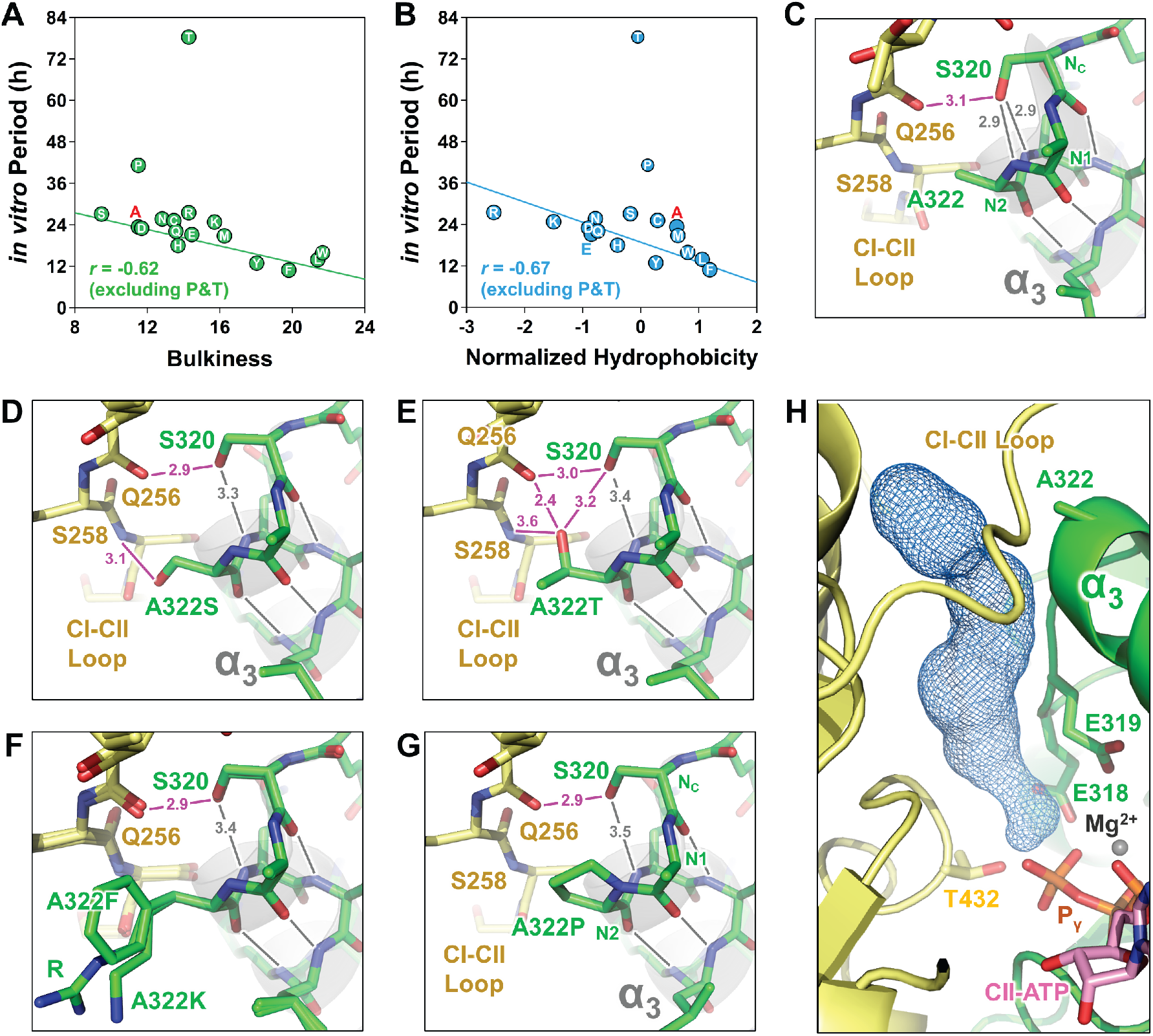
PT mechanism at the 322nd position of KaiC. (**A**) Correlation between *in vitro* periods at 30°C and bulkiness (34) of the 322nd residues. The single letter indicated at each data point represents the amino acid after substitution. Data points corresponding to KaiC^WT^ are shown in red single letters. For data points with small errors, error bars are hidden behind the markers. (**B**) Correlation between *in vitro* periods 30°C and normalized consensus hydrophobicity (35) of the 322nd residues. (**C**) A zoomed-in-view of A322, α_3_-helix (gray), and CI-CII loop (yellow) in the crystal structure of KaiC^WT^ (PDB ID: 2GBL). Nc, N1, and N2 represents a capping residue, the most N-terminal residue, and the second most N-terminal residue of the α_3_-helix, respectively (38). Gray lines indicate potential hydrogen bonds stabilizing the α_3_-helix. Magenta lines and numbers indicate other hydrogen bonds and their distances, respectively. Potential hydrogen bonds or electrostatic interactions in predicted structures of (**D**) KaiC^A322S^, (**E**) KaiC^A322T^, (**F**) KaiC^A322R^, KaiC^A322K^, and KaiC^A322F^, and (**G**) KaiC^A322P^ using AlphaFold 3 (44). (**H**) A tunnel-shaped void (navy mesh) originating from CII-ATP and extending though CII-CII interface, A322, and the CI-CII loop into the solvent (PDB ID: 2GBL).

This interpretation is also reasonable from the perspective of the physicochemical properties of the 322nd residue. Even when the parameters such as volume and bulkiness are set aside, the P-cycle frequencies still correlate somewhat with hydrophobicity or polarity of the 322nd residues. While the mutants possessing the hydrophobic 322nd residues are populated in the upper-right quadrant relative to KaiC^WT^, the lower-left quadrant is dominated by the variants harboring the polar 322nd residues (Fig. 4D). In fact, with the exception of KaiC^A322T^ and KaiC^A322P^, the P-cycle frequencies are moderately correlated with hydrophobicity measures proposed by Eisenberg et al. (35) (*r* = −0.67 in Fig. 5B). However, considering that hydrophobicity also shows a correlation with bulkiness, we conclude that the hydrophobicity dependence is merely apparent and that it is actually the modest dependence on bulkiness.

According to the crystal structure of KaiC^WT^ (21), the side-chain of A322 is not buried within the CII–CII interface. Instead, it is located in close proximity to a loop connecting the CI and CII domains (CI–CIl loop) of a neighboring protomer (Fig. 1A and yellow in Fig. 5C). These structural features provide an explanation for the variants possessing an oxygen atom at the γ position of the 322nd residue. As shown in Fig. 5D, we can construct a structural model of KaiC^A322S^ in which the oxygen atom at the γ position of S322 forms a hydrogen bond with the amide hydrogen of S258 in the CI–CIl loop without serious steric collisions with its surroundings. In a structural model of KaiC^A322T^ (Fig. 5E), the interaction with the CI–CIl loop is further strengthened by forming hydrogen-bonding and electrostatic-interaction networks among the side chain of T322, the side chain of S320, the main-chain carbonyl oxygen of Q256, and the amide hydrogen of S258. Interestingly, KaiC^A322S^ and KaiC^A322T^ both share the characteristics of having the lower ATPase activities and longer periods than KaiC^WT^. Therefore, the prolonged cycles and reduced CI-ATPase activities observed in these two mutants can be explained by restricted mobility or structural changes in the CI–CII loop of the adjacent protomer. This view is consistent with recent findings that point mutations in the CI–CIl loop have altered the period (36).

In this context, the CI–CII loop in other 322nd variants with bulkier side chains than Thr should undergo structural changes due to their steric hindrance. Nevertheless, the dependences of the period on bulkiness of the 322nd residues are not as pronounced as that of the 402nd residues (Fig. 4E and 5A). This apparent contradiction can be explained by the two structural features observed around the 322nd position. One is an intrinsically flexible nature of the CI–CII loop. Although numerous crystal structures of KaiC have been reported to date (20, 21, 31, 37), the electron density for the CI–CII loop is often unclear due to its high degree of structural freedom. Thus, it is reasonable to suggest that the CI–CII loop retains an ability to change its conformation in a flexible and passive manner in response to structural perturbations from the surrounding environment. The other is that the side chain of A322 remains partially exposed to the solvent in KaiC^WT^ (Fig. 5C). The side chains with a large volume but moderate bulkiness, such as Arg and Lys, can be exposed to the solvent by bending at the C_β_ position. In structural models of KaiC^A322R^, KaiC^A322K^, and KaiC^A322F^ (Fig. 5F), the 322nd residues are oriented toward the solvent to minimize their collisions with the CI–CII loop.

KaiC^A322P^ appears to be a rather exceptional case among the A322 variants. In KaiC^WT^, A322 is located in the N-terminal region of the α_3_-helix (Fig. 5C). The main-chain hydrogen-bonding pattern (i) > NH and O=C < (i+4) begins at S320 and its side chain caps the N-terminus of the α_3_-helix by forming two hydrogen bonds with amide hydrogens of A322 and Q323 (gray lines in Fig 5C-G). Thus, according to the previous nomenclature (38), A320 is assigned as the N-terminal capping residue (N_C_) of the α_3_-helix, R321 as the most N-terminal residue (N1), and A322 as the second most N-terminal residue (N2). Since the N2 position is in the first N-terminal turn of α-helices, even a proline residue exhibits a modest preference for the N2 position (38). On the other hand, a previous study using model peptides have reported that placing Pro at the N2 site destabilizes the α-helical structure (39). Consistently, in a structural model of KaiC^A322P^ (Fig. 5G), the capping hydrogen-bond between S320 and P322 is intrinsically absent, and that between S320 and Q323 is either disrupted or at least weakened due to the steric hinderance of the Pro residue introduced at the N2 position of the α_3_-helix. The A322P substitution may have caused a fray of the N-terminus of the α_3_-helix, indirectly affecting the structure and mobility of the CI-CII loop.

To the best of our knowledge, it is rare for the KaiC mutants with altered periods to exhibit changes not only in the CI-ATPase activity but also in the CII-ATPase activity (Fig. 4F). The N-terminus of the α_3_-helix is adjacent to dual-catalytic glutamates (E318 and E319) that constitute the CII active site. Therefore, it would not be surprising if the 322nd mutations affected the reaction site of the CII domain, but it is not necessarily obvious why the CII-ATPase increased so markedly in the short-period variants. In the crystal structure of fully-dephosphorylated KaiC (20), we found a tunnel-shaped void that runs through the CII-CII interface (Fig. 5H). This void extends from a region nearby P_γ_ of CII-ATP, passes through the CII-CII interface, narrows near A322 and the CI-CII loop, and then reaches the solvent. If the structure or mobility of the CI-CII loop is altered by the bulky 322nd residues, water molecules may be more efficiently recruited from the bulk into the CII active site through the cavity running near the 322nd position, ultimately acting as lytic water molecules that attack CII-ATP (31).

In previous studies (26-28), the P-cycle frequency has been discussed under an approximation that the total ATP consumption by KaiC is similar to the CI-ATPase activity. Since the CII-ATPase activity remained unchanged in the long-period 322nd mutants (blue bars in *SI Appendix*, Fig. S6), the approximation may not pose a significant problem for analyzing long-period mutants, as long as their periods are not extremely long. On the other hand, as seen in the short-period mutants such as KaiC^A322L^ and KaiC^A322F^, the CI-ATPase and CII-ATPase activities may increase or decrease individually, even though the combined activity increases monotonically as the period shortens (*SI Appendix*, Fig. S6). It remains unclear how the energies released in the CI and CII domains are balanced, and how this balance affects the regulation of the cycle length as well as temperature compensation. Since there are still few examples of the KaiC mutants with altered periods and elevated CII-ATPase activity, future research is needed to elucidate how the increased CII-ATPase activity contributes to cycle shortening.

## Materials and Methods

### Expression and purification of the 322nd mutants

KaiC^WT^ and its 322nd mutants were expressed in *Escherichia coli* BL21 cells as glutathione S-transferase (GST)-tagged (pGEX-6P-1) or hexa-histidine (His)-tagged (pET-3a) form and purified as described previously (40, 41). Purified KaiCs were dissolved in a Tris-buffer containing 20 mM Tris-HCl (pH 8.0), 150 mM NaCl, 0.5 mM EDTA, 1 mM ATP, 5 mM MgCl_2_, and 1mM DTT and stored at −80°C.

### *In vitro* KaiC phosphorylation cycle

KaiC (0.2 mg/ml), KaiA (0.04 mg/ml), and KaiB (0.04 mg/ml) were mixed in the Tris-buffer to reconstruct the P-cycle. An aliquot was taken from the mixture incubated at 30°C and 40°C at regular intervals. The collected samples were subjected to sodium dodecyl sulfate-polyacrylamide gel electrophoresis. The ratio of the phosphorylated KaiC was determined using a LOUPE software (42).

### ATPase measurements

The ATPase activity of KaiC (0.2–0.4 mg/ml) was measured at 30°C with a high-performance liquid chromatography system (JASCO) (43). The number of ATP molecules hydrolyzed into ADP molecules per KaiC monomer per unit time was estimated by separating accumulated ADP on an InertSustain C18 column (2 µm, 3.0 × 50 mm) (GL Sciences) equilibrated with a buffer containing 16% acetonitrile, 20 mM ammonium dihydrogenphosphate (pH 8.5), and 10 mM tetrabutylammoniumu hydrogensulfate. The flow rate was set at 0.4 ml/min.

### KaiC autodephosphorylation assay

Dephosphorylation kinetics of KaiC^WT^ and its mutants were measured at 30°C in the absence of KaiA and KaiB. Turnover rates of the unsynchronized P-cycle were calculated using a four-state model (33).

### *In vivo* bioluminescence measurements

Bioluminescence rhythms of a reporter strain of *Synechococcus elongatus* PCC 7942 were measured as described previously (19, 36). The cyanobacterial cells carrying the P*kaiBC::luxAB* reporter construct were grown on BG-11 medium under continuous light illumination (LL) at 50 µmol m^-2^ s^-1^ at 30°C for 7 d. After subjecting them to 12-h light and 12-h dark conditions, the cells were transferred to LL at 30°C and 40°C (40 µmol m^-2^ s^-1^ from an LED daylight lamp). The bioluminescence profiles were monitored by a system equipped with a photomultiplier tube.

### Structure prediction using AlphaFold 3

The amino acid sequences of KaiC^A322S^, KaiC^A322T^, KaiC^A322P^, KaiC^A322K^, KaiC^A322R^, and KaiC^A322F^ were submitted to the AlphaFold server (44). Hexameric KaiC ring structures were predicted with twelve ATP molecules and twelve Mg^2+^ ions bound at the active sites. All predictions yielded high confidence results with predicted Template Modeling (pTM) and interface predicted Template Modeling (ipTM) scores of 0.90-0.93 and ranking scores of 0.92-0.96.

## Acknowledgments

This study was supported by JSPS Grants-in-Aid for Scientific Research (22H04984 to S.A., 24H02301 to S.A., 26H01830 to Y.F., 26K02027 to K.I.-M., and 25H02446 to K.I.-M.), and partly by Takeda Science Foundation (to S.A.) and Toyoaki Scholarship Foundation (to S.A.).

## References

1. C. S. Pittendrigh, Temporal Organization - Reflections of a Darwinian Clock-Watcher. Annu. Rev. Physiol. 55, 16–54 (1993).

2. R. J. Konopka, S. Benzer, Clock Mutants of Drosophila-Melanogaster. Proc. Natl. Acad. Sci. U. S. A. 68, 2112–2116 (1971).

3. J. S. Takahashi, The 50th anniversary of the Konopka and Benzer 1971 paper in PNAS: “Clock Mutants of Drosophila melanogaster”. Proc. Natl. Acad. Sci. U. S. A. 118, e2110171118 (2021).

4. M. K. Baylies, L. B. Vosshall, A. Sehgal, M. W. Young, New short period mutations of the Drosophila clock gene per. Neuron 9, 575–581 (1992).

5. D. S. Garbe et al., Cooperative interaction between phosphorylation sites on PERIOD maintains circadian period in Drosophila. PLoS Genet 9, e1003749 (2013).

6. R. Narasimamurthy, D. M. Virshup, The phosphorylation switch that regulates ticking of the circadian clock. Mol. Cell 81, 1133–1146 (2021).

7. N. Gekakis et al., Isolation of timeless by PER protein interaction: defective interaction between timeless protein and long-period mutant PERL. Science 270, 811–815 (1995).

8. L. Saez, M. W. Young, Regulation of nuclear entry of the Drosophila clock proteins period and timeless. Neuron 17, 911–920 (1996).

9. K. L. Ode et al., Knockout-Rescue Embryonic Stem Cell-Derived Mouse Reveals Circadian-Period Control by Quality and Quantity of CRY1. Mol. Cell 65, 176–190 (2017).

10. C. L. Baker, A. N. Kettenbach, J. J. Loros, S. A. Gerber, J. C. Dunlap, Quantitative proteomics reveals a dynamic interactome and phase-specific phosphorylation in the Neurospora circadian clock. Mol. Cell 34, 354–363 (2009).

11. M. Gallego, D. M. Virshup, Post-translational modifications regulate the ticking of the circadian clock. Nat. Rev. Mol. Cell Biol. 8, 139–148 (2007).

12. P. Gao et al., Phosphorylation of the cryptochrome 1 C-terminal tail regulates circadian period length. J. Biol. Chem. 288, 35277–35286 (2013).

13. B. Wang, E. L. Stevenson, J. C. Dunlap, Functional analysis of 110 phosphorylation sites on the circadian clock protein FRQ identifies clusters determining period length and temperature compensation. G3 Genes|Genomes|Genetics 13, jkac334 (2023).

14. S. Miller et al., CRY2 isoform selectivity of a circadian clock modulator with antiglioblastoma efficacy. Proc. Natl. Acad. Sci. U. S. A. 119, e2203936119 (2022).

15. J. Landskron, K. F. Chen, E. Wolf, R. Stanewsky, A role for the PERIOD:PERIOD homodimer in the Drosophila circadian clock. PLoS Biol 7, e3 (2009).

16. O. Yildiz et al., Crystal structure and interactions of the PAS repeat region of the Drosophila clock protein PERIOD. Mol. Cell 17, 69–82 (2005).

17. K. L. Toh et al., An hPer2 phosphorylation site mutation in familial advanced sleep phase syndrome. Science 291, 1040–1043 (2001).

18. Y. Xu et al., Modeling of a human circadian mutation yields insights into clock regulation by PER2. Cell 128, 59–70 (2007).

19. M. Ishiura et al., Expression of a gene cluster kaiABC as a circadian feedback process in cyanobacteria. Science 281, 1519–1523 (1998).

20. Y. Furuike et al., Elucidation of master allostery essential for circadian clock oscillation in cyanobacteria. Sci. Adv. 8, eabm8990 (2022).

21. R. Pattanayek et al., Visualizing a circadian clock protein: Crystal structure of KaiC and functional insights. Mol. Cell 15, 375–388 (2004).

22. M. Nakajima et al., Reconstitution of circadian oscillation of cyanobacterial KaiC phosphorylation in vitro. Science 308, 414–415 (2005).

23. T. Nishiwaki et al., A sequential program of dual phosphorylation of KaiC as a basis for circadian rhythm in cyanobacteria. EMBO J. 26, 4029–4037 (2007).

24. M. J. Rust, J. S. Markson, W. S. Lane, D. S. Fisher, E. K. O’Shea, Ordered phosphorylation governs oscillation of a three-protein circadian clock. Science 318, 809–812 (2007).

25. A. Mukaiyama et al., Evolutionary origins of self-sustained Kai protein circadian oscillators in cyanobacteria. Nat. Commun. 16, 4541 (2025).

26. J. Abe et al., Atomic-scale origins of slowness in the cyanobacterial circadian clock. Science 349, 312–316 (2015).

27. K. Terauchi et al., ATPase activity of KaiC determines the basic timing for circadian clock of cyanobacteria. Proc. Natl. Acad. Sci. U. S. A. 104, 16377–16381 (2007).

28. K. Ito-Miwa, Y. Furuike, S. Akiyama, T. Kondo, Tuning the circadian period of cyanobacteria up to 6.6 days by the single amino acid substitutions in KaiC. Proc. Natl. Acad. Sci. U. S. A. 117, 20926–20931 (2020).

29. J. A. Swan et al., Coupling of distant ATPase domains in the circadian clock protein KaiC. Nat. Struct. Mol. Biol. 29, 759–766 (2022).

30. M. Elias, G. Wieczorek, S. Rosenne, D. S. Tawfik, The universality of enzymatic rate-temperature dependency. Trends Biochem. Sci. 39, 1–7 (2014).

31. Y. Furuike et al., Regulation mechanisms of the dual ATPase in KaiC. Proc. Natl. Acad. Sci. U. S. A. 119, e2119627119 (2022).

32. C. Phong, J. S. Markson, C. M. Wilhoite, M. J. Rust, Robust and tunable circadian rhythms from differentially sensitive catalytic domains. Proc. Natl. Acad. Sci. U. S. A. 110, 1124–1129 (2013).

33. Y. Furuike, Y. Onoue, S. Saito, T. Mori, S. Akiyama, The priming phosphorylation of KaiC is activated by the release of its autokinase autoinhibition. PNAS Nexus 4, pgaf136 (2025).

34. J. M. Zimmerman, N. Eliezer, R. Simha, The characterization of amino acid sequences in proteins by statistical methods. J. Theor. Biol. 21, 170–201 (1968).

35. D. Eisenberg, E. Schwarz, M. Komaromy, R. Wall, Analysis of membrane and surface protein sequences with the hydrophobic moment plot. J. Mol. Biol. 179, 125–142 (1984).

36. K. Ito-Miwa, K. Imai, K. Terauchi, T. Kondo, Intrinsic period stability of the cyanobacterial circadian oscillator across in vitro and in vivo conditions. Proc. Natl. Acad. Sci. U. S. A. 123, e2526714123 (2026).

37. R. Pattanayek et al., Structures of KaiC Circadian Clock Mutant Proteins: A New Phosphorylation Site at T426 and Mechanisms of Kinase, ATPase and Phosphatase. Plos One 4, e7529 (2009).

38. R. Aurora, G. D. Rose, Helix capping. Protein Sci. 7, 21–38 (1998).

39. D. A. E. Cochran, A. J. Doig, Effect of the N2 residue on the stability of the α-helix for all 20 amino acids. Protein Sci. 10, 1305–1311 (2001).

40. T. Nishiwaki et al., Role of KaiC phosphorylation in the circadian clock system of Synechococcus elongatus PCC 7942. Proc. Natl. Acad. Sci. U. S. A. 101, 13927–13932 (2004).

41. D. Y. Ouyang et al., Development and Optimization of Expression, Purification, and ATPase Assay of KaiC for Medium-Throughput Screening of Circadian Clock Mutants in Cyanobacteria. Int. J. Mol. Sci. 20, 2789 (2019).

42. Y. Furuike, J. Abe, A. Mukaiyama, S. Akiyama, Accelerating in vitro studies on circadian clock systems using an automated sampling device. Biophys. Physicobiol. 13, 235–241 (2016).

43. Y. Murayama et al., Tracking and visualizing the circadian ticking of the cyanobacterial clock protein KaiC in solution. EMBO J. 30, 68–78 (2011).

44. J. Abramson et al., Accurate structure prediction of biomolecular interactions with AlphaFold 3. Nature 630 (2024).

