## Supplementary material for "Identification of a Third Period-tuning Site in Cyanobacterial Clock Protein KaiC": SI Appendix

**This PDF file includes:**

Figures S1 to S7
Tables S1
SI References

### Figures

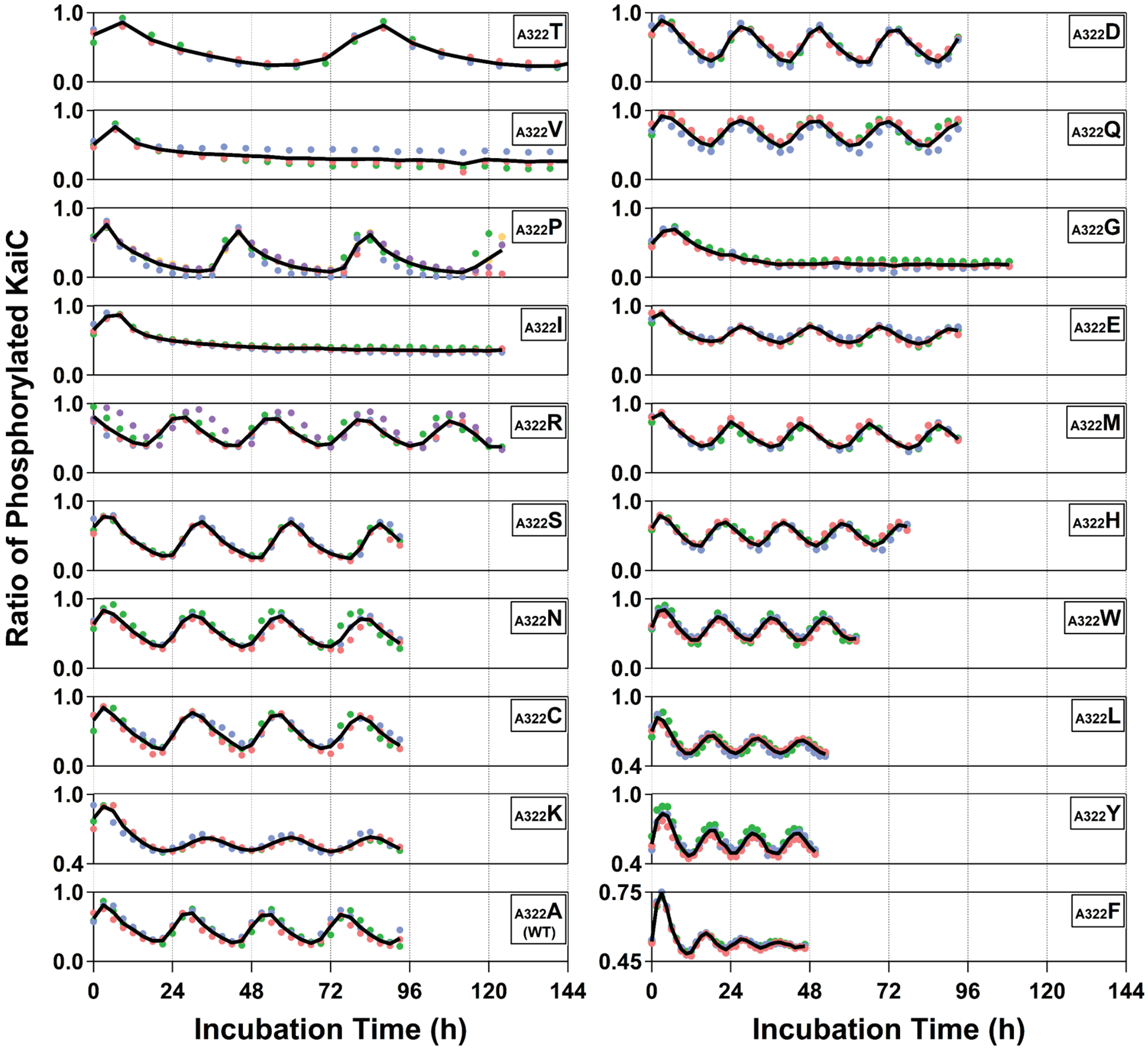


**Fig. S1.** *In vitro* phosphorylation cycles of $KaiC^{WT}$ and a series of its mutants with substitutions at the 322nd position at 30°C. Each black line corresponds to the mean from independent experiments (blue, red, green, orange, and purple circles) conducted on different days using either biological and technical replicates. Note that mutants with lower amplitudes ($KaiC^{A322K}$, $KaiC^{A322L}$, $KaiC^{A322Y}$, and $KaiC^{A322F}$) are plotted using adjusted vertical scale ranges for clarity of presentation. The samples sizes (n): n = 3 except for $KaiC^{A322P}$ (n = 5) and $KaiC^{A322R}$ (n = 5).

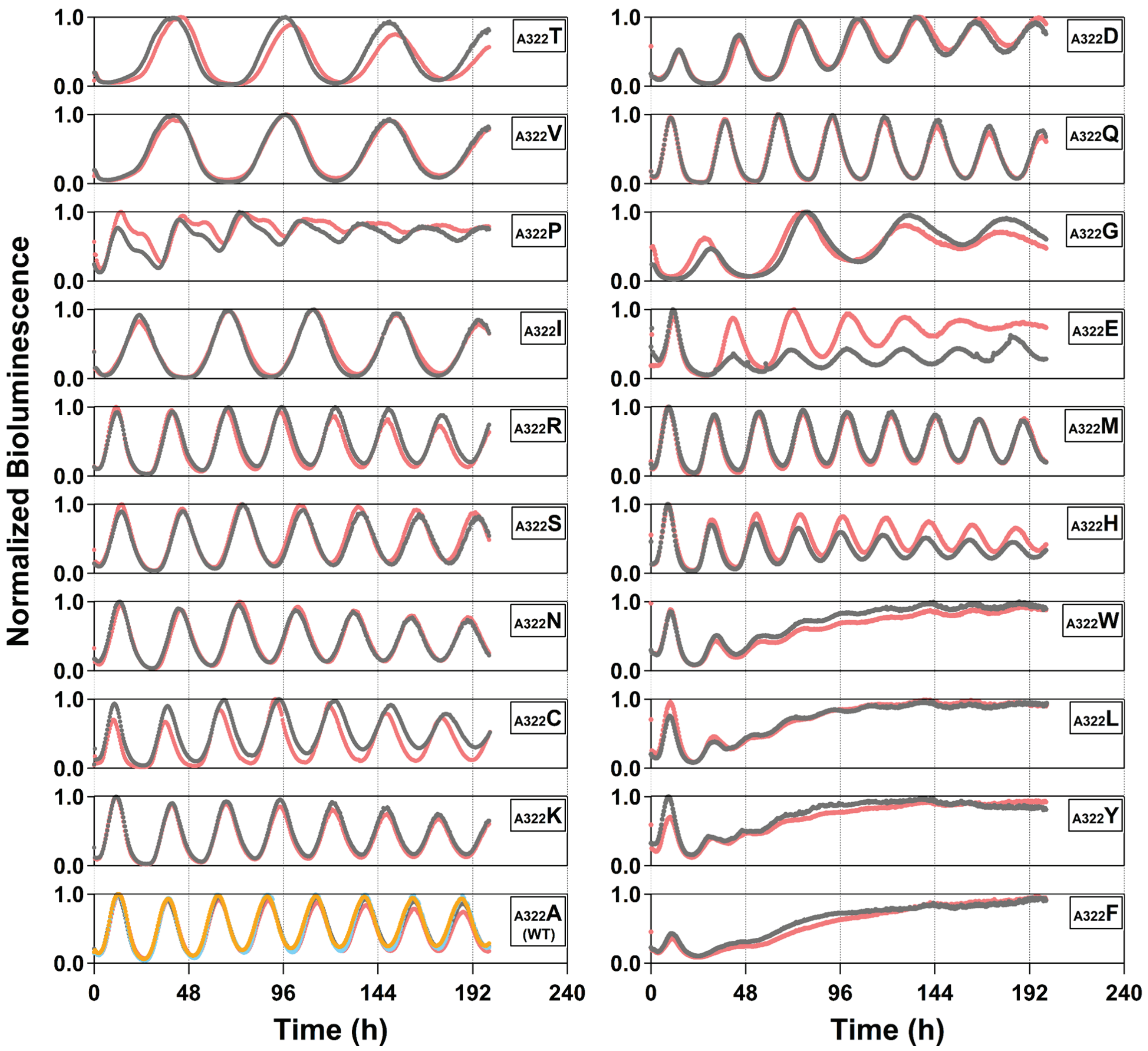


**Fig. S2.** *In vivo* bioluminescence rhythms of *Synechococcus elongatus* PCC 7942 reporter strains carrying *kaiC*$^{WT}$ and a series of its mutants with substitutions at the 322nd position at 30°C under conditions of constant light. The samples sizes (n): n = 2 except for KaiC$^{WT}$ (n = 4).

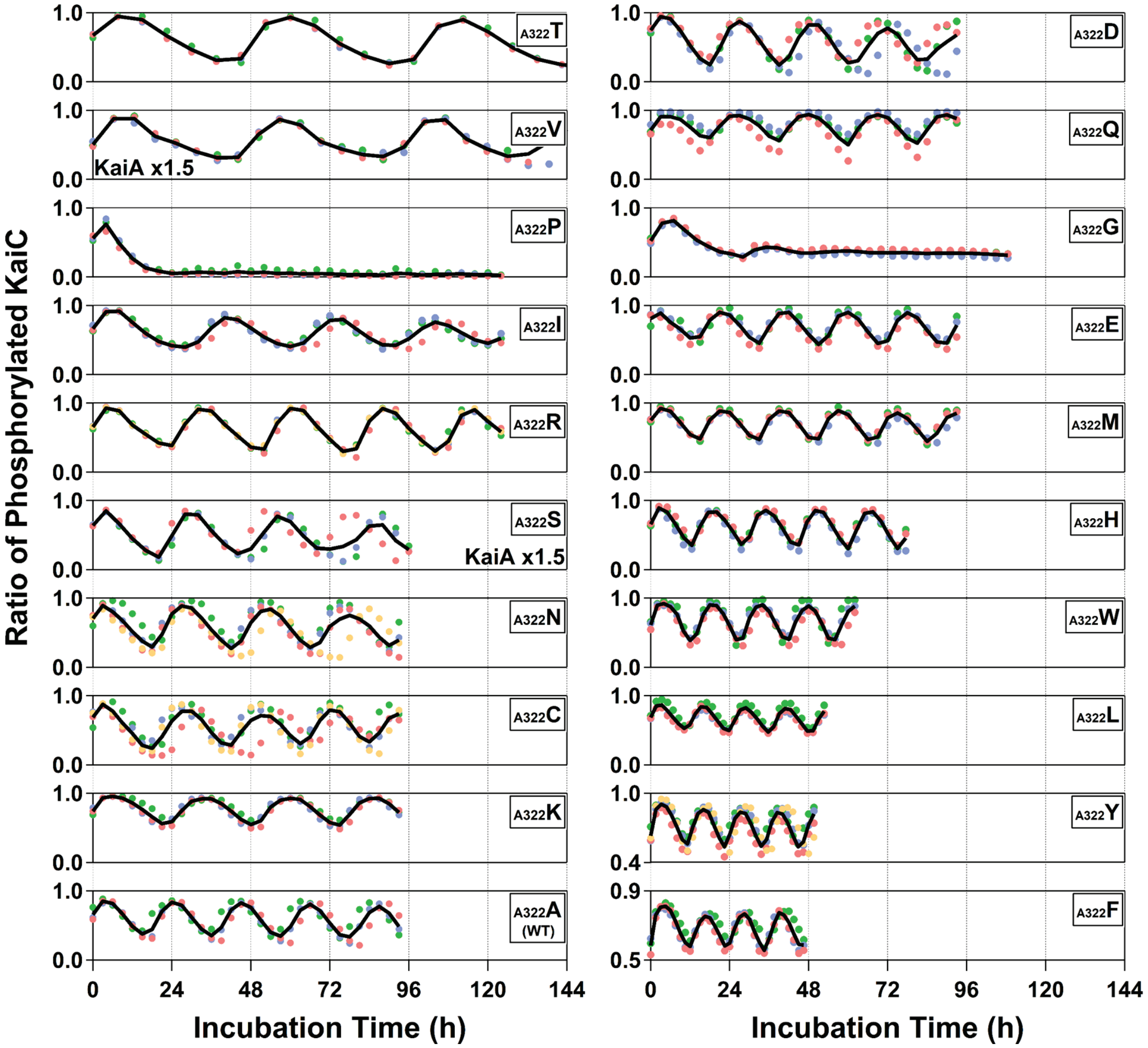


**Fig. S3.** *In vitro* phosphorylation cycles of KaiC$^{WT}$ and a series of its mutants with substitutions at the 322nd position at 40°C. Each black line corresponds to the mean from independent experiments (blue, red, green, and orange circles) conducted on different days using either biological and technical replicates. Note that mutants with lower amplitudes (KaiC$^{A322Y}$, and KaiC$^{A322F}$) are plotted using adjusted vertical scale ranges for clarity of presentation. To achieve the reproducibility of rhythms, KaiC$^{A322S}$ and KaiC$^{A322V}$ at 40°C required 1.5-fold (0.06 mg/ml) higher KaiA concentrations. The samples sizes (n): n = 3 except for KaiC$^{A322R}$ (n = 4), KaiC$^{A322N}$ (n = 4), KaiC$^{A322C}$ (n = 4), KaiC$^{A322Y}$ (n = 4).

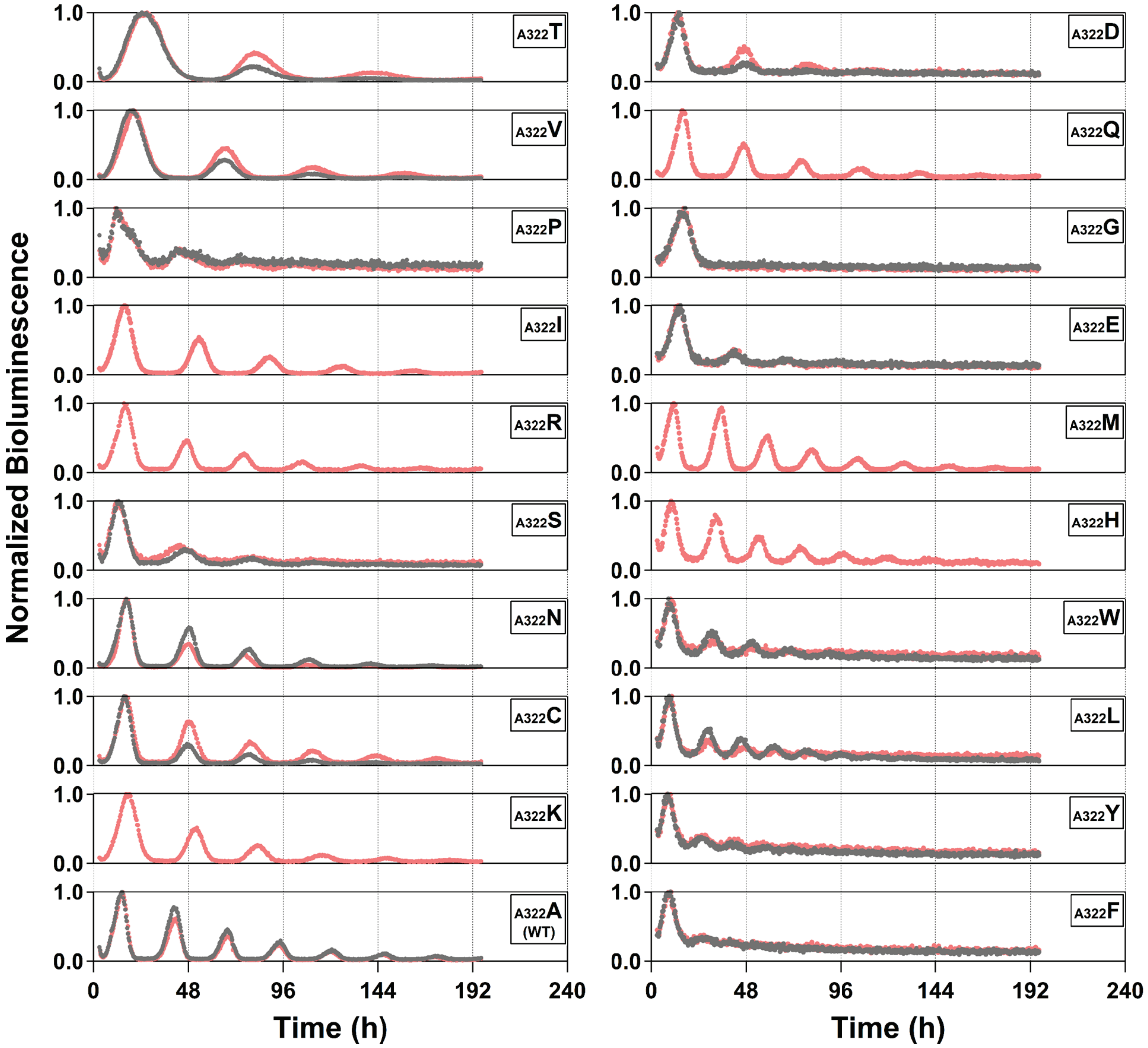


**Fig. S4.** *In vivo* bioluminescence rhythms of *Synechococcus elongatus* PCC 7942 reporter strains carrying *kaiC*$^{WT}$ and a series of its mutants with substitutions at the 322nd position at 40°C under conditions of constant light. The samples sizes (n): n = 2 except for KaiC$^{A322I}$ (n = 1), KaiC$^{A322R}$ (n = 1), KaiC$^{A322K}$ (n = 1), KaiC$^{A322Q}$ (n = 1), KaiC$^{A322M}$ (n = 1), and KaiC$^{A322H}$ (n = 1).

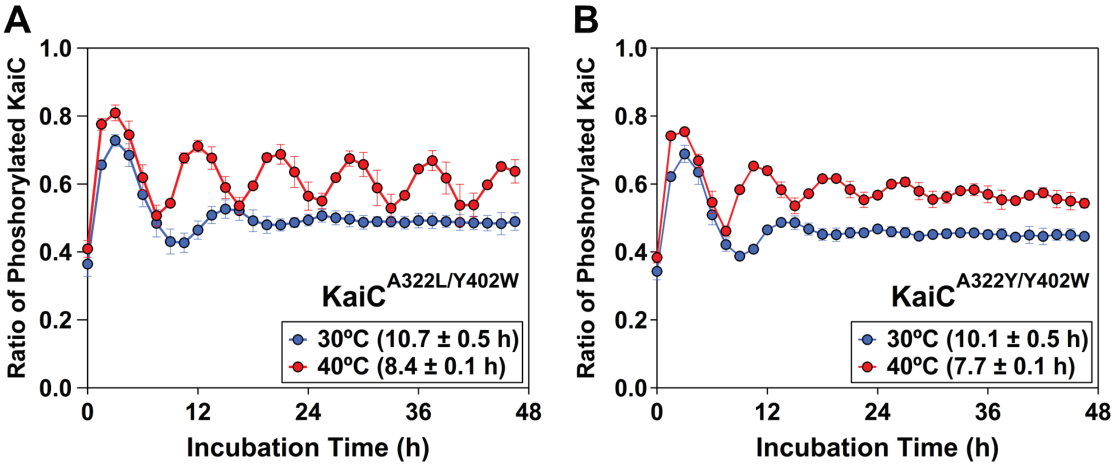


**Fig. S5.** *In vitro* phosphorylation cycles of 322nd variants with the Y402W substitution at 30°C (blue) and 40°C (red). (**A**) KaiC$^{A322L/Y402W}$. (**B**) KaiC$^{A322Y/Y402W}$. The samples sizes (n): KaiC$^{A322L/Y402W}$ (n = 3) and KaiC$^{A322Y/Y402W}$ (n = 3).

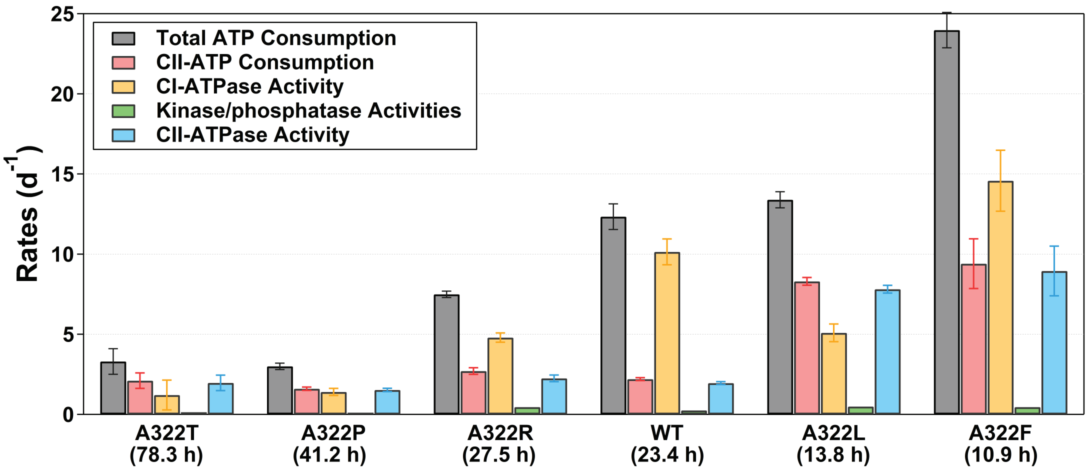


**Fig. S6.** Total ATP consumption rates (gray bars) at 30°C and their breakdown for $KaiC^{A322T}$, $KaiC^{A322P}$, $KaiC^{A322R}$, $KaiC^{WT}$, $KaiC^{A322L}$, and $KaiC^{A322F}$. Values in parenthesis represent P-cycle periods at 30°C. CII-ATP consumption rates (red bars) were estimated by introducing the E77Q/E78Q substitutions. CI-ATPase activities (orange bars) were determined as the difference in the total ATP consumption rates between the samples with and without the E77Q/E78Q substitutions. Rates of the ATP utilization as the kinase and phosphatase (green bars) were calculated using turnover rates of unsynchronized P-cycles of KaiC in the absence of KaiA and KaiB (*SI Appendix*, Fig. S7). CII-ATPase activities (blue bars) were estimated by subtracting the kinase/phosphatase activities from the CII-ATP consumption rates.

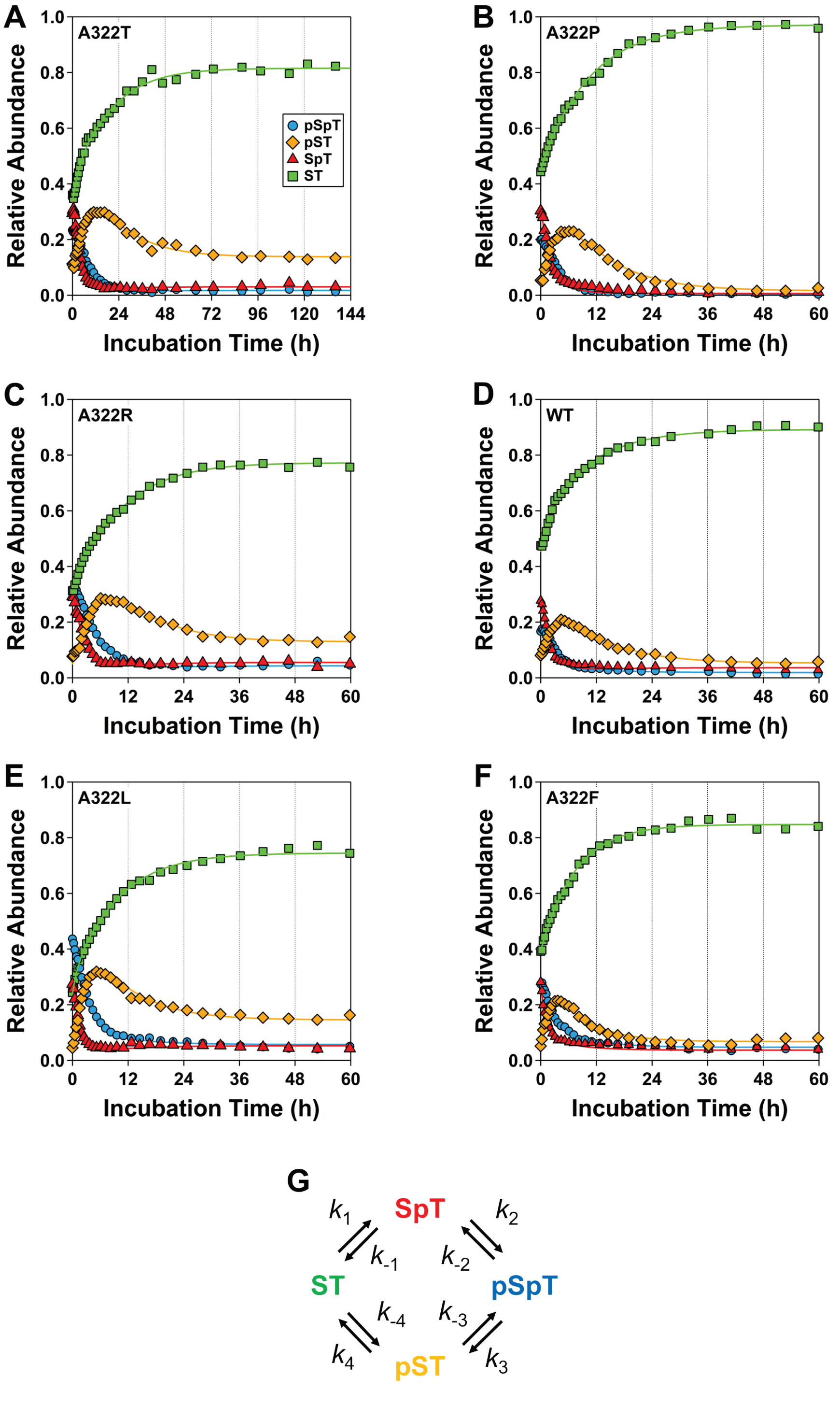


**Fig. S7.** Autodephosphorylation kinetics of (**A**) KaiC$^{A322T}$, (**B**) KaiC$^{A322P}$, (**C**) KaiC$^{A322R}$, (**D**) KaiC$^{WT}$, (**E**) KaiC$^{A322L}$, and (**F**) KaiC$^{A322F}$ at 30°C in the absence of KaiA and KaiB. ST (green squares): fully dephosphorylated KaiC. SpT (red triangles): KaiC with phosphorylated T432. pSpT (blue circles): KaiC with both S431 and T432 phosphorylated. pST (orange diamonds): KaiC with phosphorylated S431. Solid lines represent the optimal fits obtained from global nonlinear least-squares analysis using (**G**) the four-state model (1). Obtained kinetic parameters are summarized in *SI Appendix*, Table S1.

**Tables**

**Table S1.** Rate constants of the four-state model.

| Rate Constants | $KaiC^{A322T}$ | $KaiC^{A322P}$ | $KaiC^{A322R}$ | $KaiC^{WT}$ | $KaiC^{A322L}$ | $KaiC^{A322F}$ |
|---|---|---|---|---|---|---|
| $k_1$ ($h^{-1}$) | 0.009 | 0.003 | 0.029 | 0.018 | 0.041 | 0.022 |
| $k_{-1}$ ($h^{-1}$) | 0.139 | 0.189 | 0.229 | 0.308 | 0.392 | 0.260 |
| $k_2$ ($h^{-1}$) | 0.096 | 0.374 | 0.201 | 0.152 | 0.209 | 0.833 |
| $k_{-2}$ ($h^{-1}$) | 0.000 | 0.160 | 0.038 | 0.015 | 0.013 | 0.448 |
| $k_3$ ($h^{-1}$) | 0.168 | 0.436 | 0.220 | 0.387 | 0.312 | 0.458 |
| $k_{-3}$ ($h^{-1}$) | 0.000 | 0.019 | 0 | 0.038 | 0.052 | 0.181 |
| $k_4$ ($h^{-1}$) | 0.046 | 0.100 | 0.085 | 0.099 | 0.100 | 0.140 |
| $k_{-4}$ ($h^{-1}$) | 0.004 | 0 | 0.002 | 0 | 0.006 | 0 |

### SI References

1. Y. Furuike, Y. Onoue, S. Saito, T. Mori, S. Akiyama, The priming phosphorylation of KaiC is activated by the release of its autokinase autoinhibition. *PNAS Nexus* **4**, pgaf136 (2025).
